# Elevated substitution rates and increased purifying selection associated with thermophily in *Chaetomiaceae* fungi

**DOI:** 10.64898/2026.07.21.739749

**Authors:** Noah Hensen, Thibault Caron, Markus Hiltunen Thorén, Hanna Johanneson

## Abstract

Understanding the genomic consequences of thermal adaptation in fungi is crucial, as rising global temperatures are expected to have negative impacts on food safety and human health. The family *Chaetomiaceae* contains a large number of thermophilic fungal taxa, but previous studies have reported inconsistent optimal growth temperatures (OGT) for the same strains, obtained with various laboratory methods. Here we applied a standardized laboratory approach to measure OGT across strains of 17 *Chaetomiaceae* species and used a phylogenomic approach to test associations between OGT, rates of genome evolution, and strength of purifying selection. Compared to mesophiles, thermophilic fungi showed faster nucleotide substitution rates. In addition, thermophiles showed lower dN/dS ratios than mesophiles, suggesting stronger purifying selection on conserved orthologs. We hypothesize that the elevated substitution rates are linked to high growth rates, as thermophilic fungi grew significantly faster than mesophilic ones. Our results show that selective pressures may act at different temperatures for distinct genomic characteristics. Genome size was lower at OGT ≥ 35°C compared to mesophilic species, while GC content did not show a large difference between mesophiles and thermotolerant species, but increased in thermophilic species with an OGT ≥ 45°C.

## 1. Introduction

As global temperatures rise, many species face novel environmental pressures, resulting in geographic range shifts, adaptation or risk of extinction (Bell & Collins, 2008). At the same time, the ongoing loss of those species unable to survive changing environments allows the spread of species better-adapted to the novel conditions (Turvey & Crees, 2019). Directed evolution experiments have shown that fungal species are able to overcome temperature barriers quickly (de Crecy et al., 2009), and global warming is already affecting their geographic range, abundance, and thermotolerance (Bebber et al., 2013; Fisher et al., 2012; McLean et al., 2005; Robert et al., 2015; Velásquez et al., 2018). As a result, fungi are projected to have increasingly large negative effects on food safety and human health (Ipcc, 2022), highlighting the major need for understanding the genomic basis of thermal adaptation in the fungal kingdom (Korfanty et al., 2023), including the relationship between evolutionary rates, strength of selection and adaptation to high optimal growth temperatures.

Temperature strongly influences evolutionary dynamics, in part through its effect on mutation rates, a key parameter of evolutionary potential (Chu et al., 2018). Mutations generally tend to accumulate faster at higher temperatures (Matsuba et al., 2013). However, mutation rates and substitution rates in thermophilic prokaryotes tend to be lower than in their mesophilic relatives, despite faster growth. This reflects strong selection for genomic stability, which could be contributed to, for example, more efficient DNA repair mechanisms or efficient purifying selection (Drake, 2009; Friedman et al., 2004; Jacobs & Grogan, 1997; Mackwan et al., 2008; Sabath et al., 2013; Swami, 2009; Xu et al., 2021). In addition, thermophilic prokaryotes have been shown to contain smaller genomes with higher GC content and more pronounced codon usage biases, which together enhance DNA stability and energy efficiency at high temperatures (Borisova et al., 1993; Giovannoni et al., 2014; Hu et al., 2022; Sabath et al., 2013; Yakovchuk, 2006).

Although detailed research is lacking, it has been hypothesized that thermophilic fungi show similar correlations between genomic features and high optimal growth temperatures as bacteria (Hensen et al., 2023; Steindorff et al., 2024; van Noort et al., 2013). Within the ascomycete family *Chaetomiceae*, thermophilic and thermotolerant taxa are common, and polyphyletic (Geydan et al., 2012; van den Brink et al., 2015). Beyond their utility as a model for studying thermal adaptation, *Chaetomiaceae* species are also of applied interest: many produce thermostable enzymes used in industrial processes, and the family includes opportunistic human pathogens such as *Madurella* (Van Belkum et al., 2013; van den Brink et al., 2015). Earlier research showed reduced genome sizes and increased GC contents in thermophilic *Chaetomiaceae*, compared to closely-related mesophiles (van Noort et al., 2013). Similar trends were seen in other *Chaetomiaceae* genomes compared to those of its sister family *Podosporaceae* (Hensen et al., 2023). The *Chaetomiaceae* family is further described to show high levels of codon usage bias and genome-wide selection on codon usage bias, which could confer an adaptation to higher optimal growth temperatures (OGT) by enhancing protein stability (Hensen et al., 2023; Steindorff et al., 2024). However, correlating OGT with genomic properties has remained difficult due to varying experimental methods and conflicting reports of OGT for the same fungal strains (Steindorff et al., 2024; van den Brink et al., 2015).

Despite increasing interest in fungal thermal adaptation, inconsistencies in reported growth optima and limited comparative genomic data have hindered a clear understanding of how temperature shapes fungal genome evolution. In this study we applied a standardized framework to assess OGT across 19 *Chaetomiaceae* species and integrated OGT data with phylogenetic and comparative genomic analyses. Our main aim was to assess whether evolutionary rates and strength of purifying selection on conserved single copy orthologs differ between thermophilic, thermotolerant, and mesophilic species. Additionally, we tested whether OGT correlates with genomic traits such as genome size, gene content, and measures of codon usage bias. Together, these analyses provide a comparative framework for understanding the genomic basis of thermal adaptation in fungi.

## 2. Methods

In this study, we investigated a total of 25 strains, including 23 strains of *Chaetomiaceae* species and strains of two outgroup species (*Cercophora caudata* from *Lasiosphaeriaceae*, and *Podospora didyma* from *Neoschizotheciaceae*). Full genome sequences for all species are publicly available (see Table 1 for corresponding references). Quality of genomes was assessed using BUSCO V5.3.1 with augustus species training and the sordariomycetes_odb10 database (Kriventseva et al., 2019; Manni et al., 2021; Stanke et al., 2006). All genomes had >70% completeness for single copy BUSCO genes, and 23 out of 25 assessed strains had >90% completeness for single copy BUSCO genes (Table 1), indicating high quality genomes.

**Table 1:** Fungal strains used in this study, BUSCO completeness of the genomes, and overview of assessed optimal growth temperature groups in different studies.

| Species | Strain ID | Assessed OGT |  |  | Genome publication | Single copy BUSCO genes (%) |
| --- | --- | --- | --- | --- | --- | --- |
|  |  | This study | Steindorf et al., 2024 | van den Brink et al., 2015 |  |  |
| *Achaetomium macrosporum* | CBS 532.94 |  | M | M* | Hensen et al., 2023 | 98.8 |
| *Achaetomium strumarium* | CBS 333.67 | TT | TT | T | Hensen et al., 2023 | 98.8 |
| *Canariomyces arenarius* | CBS 508.74 | TT | T | TT | Hensen et al., 2023 | 99.2 |
| *Cercophora caudata* (outgroup) | CBS 606.72 | M |  |  | Hensen et al., 2023 | 98.6 |
| *Chaetomium cochliodes* | NA |  |  |  | Zamocky et al. 2016 | 84.6 |
| *Chaetomium fimeti* | CBS 168.71 | M | M |  | Hensen et al., 2023 | 98.8 |
| *Chaetomium globulosum* | CBS 148.51 | M | M |  | Cuomo et al., 2015 | 92 |
| *Chaetomidium leptoderma* | CBS 538.74 | M | T |  | Amlacher et al. 2011 | 99 |
| *Chaetomium thermophilum* | CBS 144.50 | T |  | T | Hensen et al., 2023 | 96.3 |
| *Corynascella inaequalis* | CBS 284.82 | M | M |  | Hensen et al., 2023 | 99.1 |
| *Corynascus novoguineensis* | CBS 359.72 | TT | M |  | Berka et al. 2011 | 99.2 |
| *Corynascus sepedonium* | CBS 632.67 | TT | TT |  | Morgenstern et al. 2012 | 99.3 |
| *Crassiacarpon hotsonii* | CBS 398.93 |  | T | T* | Morgenstern et al. 2012 | 97.4 |
| *Crassiacarpon thermophilum* | CBS 405.69 | TT | T |  | Hensen et al., 2023 | 92.5 |
| *Dichotomopilus funicola* | CBS 141.50 | M |  |  | Smit et al. 2016 | 98.8 |
| *Madurella mycetomatis* | NA |  |  |  | Morgenstern et al. 2012 | 79.8 |
| *Mycothermus thermophilus* | CBS 625.91 | TT | T | T | Hensen et al., 2023 | 97.5 |
| *Parathielavia appendiculata* | CBS 731.68 | TT | M | TT | Hensen et al., 2023 | 99.1 |
| *Parathielavia hyrcaniae* | CBS 757.83 | M | M | M | Hensen et al., 2023 | 98.9 |
| *Podospora didyma* (outgroup) | CBS 232.78 | M |  |  | Hensen et al., 2023 | 98.4 |
| *Staphylotrichum longicollum* | CBS 103.79 | M | T |  | Hutchinson et al. 2016 | 97.7 |
| *Thermothelomyces heterothallica* | CBS 202.75 | T | T | T | Berka et al. 2011 | 98.8 |
| *Thermothielavioides terrestris* | NA |  |  |  | Berka et al. 2011 | 99.1 |
| *Thermothelomyces thermophila* | NA |  |  |  | Hensen et al., 2023 | 99 |
| *Trichocladium antarcticum* | CBS 123565 | M | M | M | Cuomo et al., 2015 | 98.3 |

*Classification of species into mesophilic (M), thermotolerant (TT), and thermophilic (T) species.*
*\* These assessments were made on different strains of the same species. All species belong to the Chaetomiaceae* *family, with the exception of two outgroup species, indicated with (outgroup) in the species column. The outgroup* *species Cercophora caudata and Podospora didyma belong the families Lasiosphaeriaceae and Neoschizotheciaceae,* *respectively.*

### 2.1 Optimal growth temperature and growth rate assessment

Different definitions for fungal thermophilia exist (see e.g. Morgenstern et al., 2011; Steindorff et al., 2025; van den Brink et al., 2015), and in this study we broadly follow the categories defined by van den Brink et al. (2015). Based on relative growth performance, species with the highest growth rate at ≤ 30°C were classified as mesophilic, those growing optimally at 35 - 40°C as thermotolerant, and species with an OGT ≥ 45°C as thermophilic (van den Brink et al., 2015).

We assessed optimal growth temperature (OGT) and growth rates of strains from 19 species, including both outgroup strains and 17 *Chaetomiaceae* strains (Table 1). The OGT of two additional species (*Achaetomium macrosporum* and *Thermothelomyces myriococcoides*) were directly obtained from van den Brink et al. (2015). Freeze-dried cultures of the 19 available strains were obtained from the Westerdijk Fungal Biodiversity Institute, and revived according to their instructions (Westerdijk, 2025). All cultures were maintained on 40% malt extract agar (MEA) at 20°C, with the exception of *Chaetomium thermophilum* and *Crassicarpon thermophilum*, which showed no growth at 20°C and were instead grown at 30°C. Assessments were standardized based on laboratory methods described by van den Brink et al. (2015). For inoculations, a small agar plug (roughly 3 mm in diameter) containing mycelium from the edge of a growing colony was transferred to the middle of 9 cm Petri dishes containing roughly 18 mL 4% MEA. Cultures were incubated in the dark for six days, in enclosed plastic boxes containing a 300mL beaker of water to prevent them from drying out. Temperatures ranged from 15 to 55°C with 5°C intervals.

After six days, colony size was measured. OGT was defined as the temperature at which the colony diameter was largest at the end of the incubation period. For five strains that had reached the edges of the Petri dish at day six at multiple temperatures, growth was re-analyzed by measuring the diameter of the colony on day three. Growth rates were determined by measuring the diameter of colonies at the end of inoculation compared to diameter at inoculation (3 mm). All assessments were done in triplicate.

### 2.2 Phylogenetic reconstruction

To obtain a reference phylogeny, the obtained single-copy BUSCO results were parsed to obtain genes found in all genomes in the dataset (i.e. 100% taxonomic coverage). The resulting dataset contained 2096 out of the 3817 genes in the dataset. Nucleotide sequences were aligned with MACSE V2.07, using the standard translation code (Elzanowski & Ostell, 2019; Ranwez et al., 2018). Resulting alignments were individually trimmed using TrimAl V1.4 with the default settings of the “gappy-out” parameter (Capella-Gutierrez et al., 2009). A subset of 40 randomly selected gene alignments was visually examined and showed no unambiguously aligned sequences. Single gene trees were inferred for each gene using IQ-TREE v2.1.4_beta with 1000 ultrafast bootstrap replicates, and with the -m TEST --runs 10 option to identify the best-fitting substitution model (Minh et al., 2020). For species-tree inference, a concatenation-based tree was reconstructed under the best-fit model (TIM3+F+I+G4) and evaluated with 1,000 ultrafast bootstrap replicates. In parallel, a coalescence-based species tree was generated using ASTRAL-III from gene trees inferred individually under their own best-fit models and assessed with posterior probabilities (Zhang et al., 2018). Gene and site concordance factors were estimated using IQ-TREE v2.4 with the -gcf option to assess gene concordance factors, and the maximum likelihood-based method (--scfl) for site concordance factors (Mo et al., 2023). The concatenation- and coalescence-based methods created two similar, yet distinct, trees that were both used for further substitution rate analysis.

While thermophily has been shown to be a polyphyletic trait within *Chaetomiaceae*, it is possible that the trait has evolved by horizontal transfer of genes promoting growth at a high temperature optimum. If so, trees generated from such genes would cluster species according to their OGT. We analysed phylogenetic signal of OGT on all individual gene trees and the species tree. For practical reasons, only the concatenation-based species tree was used for inference of phylogenetic signal on the species tree. Due to the discrete trait values (M, TT, T), phylogenetic signal was assessed with the M-statistic, which estimates phylogenetic signal by comparing distances from phylogenies and traits. The M-statistic ranges from 0-1 for discrete traits, with higher phylogenetic signal leading to values closer to 1. The phylogenetic signal of a trait is assumed to be significant when p < 0.05. The M-statistic was computed in R V4.6.0 with R-package phylosignalDB (R Core Team, 2022; Yao & Yuan, 2025).

Internal support of branches in trees with significant phylogenetic signal were individually examined to determine support of OGT clusters. We examined whether the number of gene trees with phylogenetic signal was higher than under randomization of OGT by creating 1000 permutations with randomized OGT across taxa. Putative functions of genes showing phylogenetic signal for OGT were inferred as follows. We first deleted alignment-gaps from the fasta files, and next annotated the gap-free versions of all 2096 BUSCO genes using InterProScan v.575-106.0 (Jones et al., 2014). Next, GOATOOLS V1.2.3 was used to search for significantly enriched GO terms in the subset of genes showing phylogenetic signal for OGT (Klopfenstein et al., 2018).

### 2.3 Substitution rate analysis

Nucleotide and amino acid substitution rates, together with the strength of purifying selection, were estimated with PAML (Álvarez-Carretero et al., 2023). More specifically, BASEML was used to analyze different models of nucleotide substitutions. The rate of nonsynonymous substitutions (dN), the rate of synonymous substitutions (dS), and the nonsynonymous/synonymous substitution rate ratio (dN/dS) were analyzed with CODEML. Here, the dN/dS ratio of protein-coding genes was used as an estimate for strength of selection during amino acid evolution. Specifically, dN/dS ratios lower than 1 indicate purifying selection and genome-wide evolutionary constraints.

For the input phylogeny, we pruned the concatenation and coalescence-based trees to exclude species without measured OGT. Trees were used unrooted, or were rooted with *Podospora didyma* when required using the R-package ggtree v3.12.0 (Yu et al., 2017, 2018). All 2096 one-to-one BUSCO genes were filtered to obtain only high-quality coding DNA sequences (CDS). These were selected based on the following criteria: the spliced CDS 1) had a length of ≥ 100 bp, 2) contained no partial codons, 3) had no internal stop codons, and 4) contained both start and stop codons. If an ortholog from any species did not fulfill these criteria, the gene was fully removed from the dataset (i.e., no missing data was allowed). Of the 2096 BUSCO genes present in all genomes in the dataset, 1162 remained after gene filtering and were used for further analysis. All 1:1 orthologous genes were aligned with PRANK, as described by van der Lee et al., 2017 (van der Lee et al., 2017). Alignment filtering was performed with Guidance2, using the default settings (guidance.pl - -program GUIDANCE - -seqType nuc - -msaProgram PRANK - -MSA Param “\+F \-codon”; v1.5) (Sela et al., 2015). The alignments filtered by Guidance2 were used for subsequent analysis.

In PAML, different models were utilized to assess global, local, and free substitution rates. The clock = 1 (BASEML) or M0N0 (CODEML) model represents a model with one global rate of nucleotide substitution or selection strength for all branches. Free substitution rates, assessed with the clock = 0 or M1N0 models, allow rates to vary freely among each branch. Local substitution rates were obtained with clock = 2 or M2N0. For the local models, we analyzed both two-way (thermophiles vs. non-thermophiles) and three-way (mesophiles, thermotolerant, and thermophiles) splits. Furthermore, branches in the local model were categorized based on OGT using both classification alternatives of van den Brink et al. (2015) and classifications obtained in this study, yielding a total of four different branch categories. Lastly, the rate of nonsynonymous substitutions and the rate of synonymous substitutions were calculated for the two-way and three-way local models and the global model. The rate of nonsynonymous substitutions was calculated by summing up the number of nonsynonymous substitutions along the tree for each of the three groups and dividing it by the total number of possible nonsynonymous substitutions along those branches. The same was done for the rate of synonymous substitutions. To verify which of the models best fit the data, likelihood ratio tests (LRTs) were performed by comparing twice the difference in log likelihood values between two models using a Chi2 distribution.

### 2.4 Genomic traits and statistics

We evaluated various genomic features and correlated them with our OGT and growth rate assessments. We obtained the following genomic variables for the included genomes from openly available data: GC content, effective number of codons (ENC), genome size, and gene number (Hensen et al., 2023, 2025; Cuomo et al., 2015; Table S1). GC content of coding sequences and GC content of the third codon position were obtained from the PAML output.

For all genomic traits, normality of data and homogeneity of variances were tested with QQplots, Shapiro-Wilk tests and Bartlett’s test, and showed no deviation from normality or heterogeneity of variances. Correlations between variables were assessed with Pearson correlation coefficients. All statistics were obtained with R version 4.4.1 in R-Studio (Rstudio Team, 2022). Results were plotted with ggplot2 (R Core Team, 2022; Wickham, 2016). Due to limited sample sizes, differences between groups (mesophilic, thermotolerant, and thermophilic strains) are only described quantitatively.

## 3. Results

### 3.1 Standardized tests clarify optimal growth temperatures within *Chaetomiaceae*

The *Chaetomiaceae* cultures showed a wide range of OGT and growth rates. Growth rates were slowest in the outgroup strain *Podospora didyma*, with a maximum growth of 15 mm after 6 days of incubation at OGT (20°C), to a growth of 80 mm within three days at 45°C in the thermophilic *Thermothelomyces heterothallica* (Table S1). For cultures from five strains (*Crassicarpon thermophilum, Thermothelomyces heterothallica, Achaetomium strumarium, Chaetomium thermophilum, Chaetomium globulosum*), the mycelium had filled the entire 9 cm Petri dish at day 6 at multiple temperatures. These were reassessed with a three-day incubation period to obtain the OGT. In a further two strains (*Canariomyces arenarius* and *Thermothelomyces heterothallica*), we found 100% relative growth at multiple temperatures (Table S1); in these cases, the upper temperature was used as OGT for genomic analysis. Eight out of 19 assessed strains could grow at 45°C. Of these, the lower end of the growth range was largely variable, from ≤15°C for *Corynascus sepedonium* to a minimal growth temperature of 30°C for *Crassicarpon thermophilum* (Figure 1). For most strains analyzed in multiple datasets, results between studies were consistent and we could confidently assign classifications (Table 1). Despite similar laboratory methods between van den Brink (2015) and our study, two inconsistencies remained. Specifically, *Achaetomium strumarium* and *Mycothermus thermophilus* were both classified as thermotolerant species in our study, while being classified as thermophilic in van den Brink et al. (2015). Therefore, further analyses of the relationship of OGT and rates of genome evolution were performed using both sets of classifications (Table 1).

**Figure 1:**
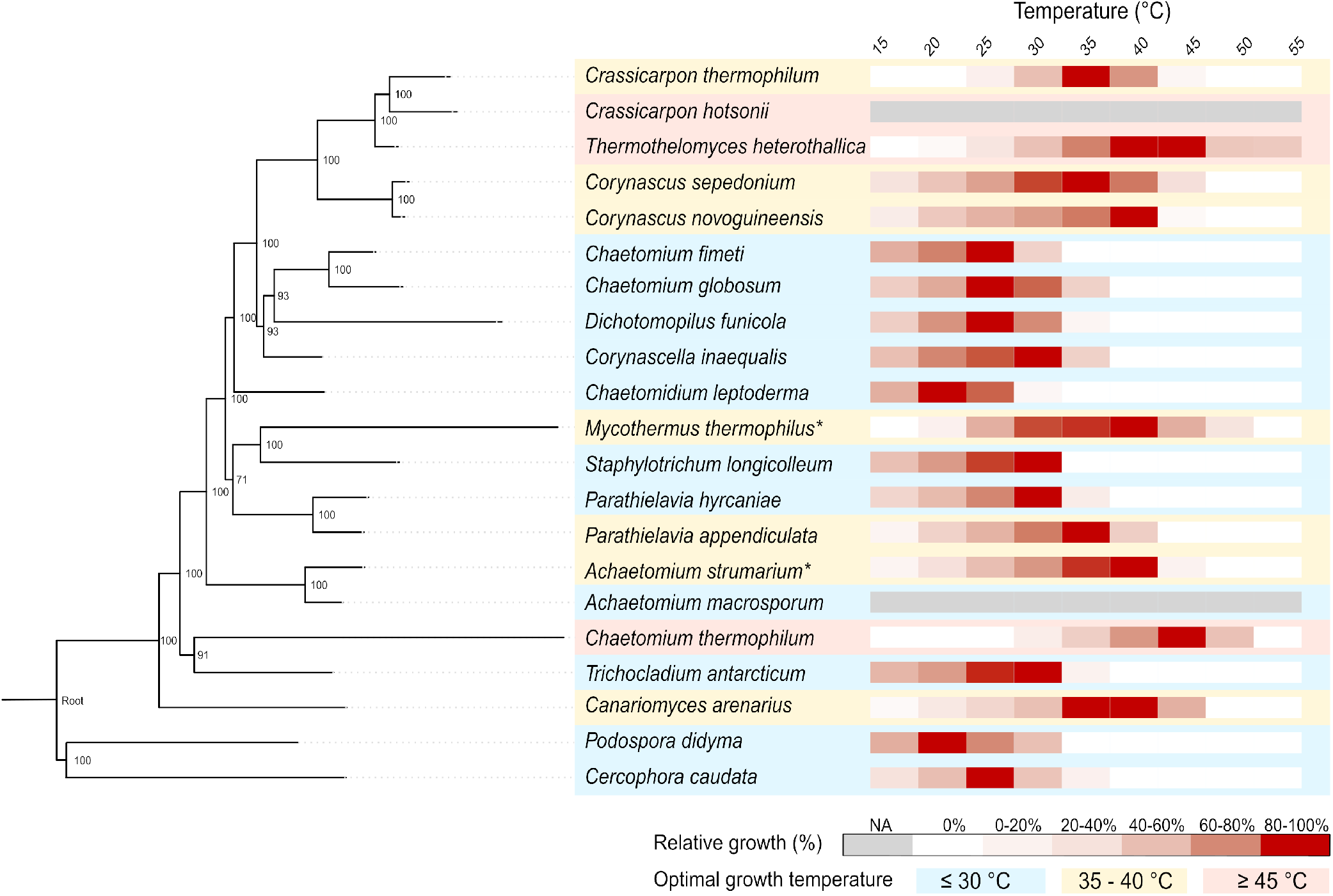
Concatenation-based tree with heatmap of relative growth rate (%) at different temperatures (°C). Maximum value is 100% growth at the OGT. Different columns show different temperatures from 15-55°C. Grey bars indicate species with known OGT classification from research by van den Brink et al., (2015), but not assessed during this project. Branches show UFBS support values. * These two species were assessed as thermophilic by van den Brink et al., (2015).

### 3.2 Evolution of thermophilia in *Chaetomiaceae*

Previous research has shown that, despite the use of high-quality data, underlying gene tree discordance remained prevalent within the *Chaetomiaceae* (Hensen et al., 2023), making it difficult to determine the exact species tree. We re-created the phylogeny of *Chaetomiaceae* with one additional taxon added to the dataset used in Hensen et al., (2023): *Chaetomium globulosum* (Table 1). Both a coalescence and a concatenation-based approach were used. While the majority of nodes were congruent between the two trees, underlying gene tree discordance remained prevalent both within and between clades, as indicated by low gene concordance factors in some branches (Figure S1). We performed subsequent substitution rate analysis on both the concatenation and the coalescence-based phylogeny and observed no difference in outcome of either general results or levels of significance. As a result, systematic differences in phylogenies between the two methods are unlikely to bias our presented results. For further analysis, we present results with the concatenation-based phylogeny (Figure 1), results for other analysis can be found in the online supplementary data repository.

We investigated if any gene trees grouped species according to their OGT, as would be expected under horizontal gene transfer within *Chaetomiaceae*. Of the 2,096 single-gene trees analyzed, 30 trees showed significant phylogenetic signal for OGT (p < 0.05) and high strength of signal according to the M statistic. Within these 30 genes, the average signal strength was 0.76 +-0.03, on a scale of 0-1. An additional 49 genes showed moderate signal (0.05 ≤ p ≤ 0.1), and lower average signal strength at 0.71 +-0.02. Most genes (2001 genes), as well as the species tree, showed no significant phylogenetic signal according to the M-statistic, and lower overall signal strengths. The average signal strength in genes without signal was 0.58 +-0.06, with the species tree having a signal strength of 0.58 and a p-value of 0.5 (Figure S2). A test with 1000 permutations of the OGT across the tree indicated that the number of trees with significant phylogenetic signal was not higher than expected under randomization of OGT (figure S3). We individually examined gene trees that showed significant phylogenetic signal and found a mix of highly supported and poorly supported branches (Figure S4). Significant enrichment of 10 GO terms was found in the subset of genes with phylogenetic signal compared to the whole set of BUSCO genes (p-value after Bonferroni correction < 0.05). Enriched genes were mainly associated with (intracellular) organelles (GO:0043299, GO:0043226, GO:0043232, GO:0043228) (Figure S5). Taken together, while some genes trees clustered species according to their trait value (M, TT, T), this number was not higher than expected by random chance and we conclude it is unlikely that thermophily has evolved by horizontal gene transfer.

### 3.3 Increased nucleotide substitution rates in thermophilic fungi

To determine if OGT is correlated with nucleotide substitution rates, we compared the fit of our data to three different substitution rate models, i.e. a global model, a free-ratio model and a local branch model (Table 2). Local models were obtained both with OGT from our laboratory assessments and from van den Brink et al (2015), no differences were observed in the results between the two classifications (Table S2). We here present results based on OGT classifications obtained in this study. The two-way split local rate model fit the data significantly better than the global rate model (Table S2), with thermophilic fungi showing average substitution rates 1.5-fold higher than mesophilic taxa (Table 2). The three-way split model gave a statistically better fit than the global model or a local model which distinguished only thermophilic versus non-thermophilic species. In the three-way split model, thermotolerant taxa showed slightly higher substitution rates than mesophilic species (1.3 vs 1.0 on average), but lower than thermophilic species (1.7).

**Table 2:** Summary statistics and parameter estimates of nucleotide substitution rates among branches in the *Chaetomiaceae*.

| Nucleotide substitution model | $\ell$ | p-value* | nucleotide substitution rates | | |
| --- | --- | --- | --- | --- | --- |
| 1, Global substitution rate | -20780796.7 | - | 1.42 (global) |  |  |
| 0, Free substitution rates | -20586861.6 | <0.01<sup>c, d, e</sup> | branch-specific rate |  |  |
| 2<sup>a</sup>, Local model | -20763126.2 | <0.01<sup>c, e</sup> | 1.00 (NT) | 1.51 (T) |  |
| 2<sup>b</sup>, Local model | -20751267.8 | <0.01<sup>c</sup> | 1.00 (M) | 1.66 (T) | 1.29 (TT) |

1) Global substitution rate, assessed with the clock = 1 model.
0) Free substitution rates along each branch of the phylogeny, assessed with the clock = 0 model.
2<sup>a</sup>) Local substitution rate with two groups, non-thermophiles (NT) and thermophiles (T), assessed with the clock = 2 model.
2<sup>b</sup>) Local substitution rate with three-way split between mesophiles (M), thermotolerant (TT), and thermophiles (T), assessed with the clock = 2 model.
<sup>c</sup>) The null hypothesis is the global model (model clock = 1) in the likelihood ratio tests (LRT).
<sup>d</sup>) The null hypothesis is the 2<sup>a</sup> local model in the LRT.
<sup>e</sup>) The null hypothesis is the 2<sup>b</sup> local model in the LRT.
\* Significant if the alternative hypothesis is a significantly better fit than null hypothesis. Full statistics and additional assessments are shown in table S2 and in the online data repository.

The free-ratio model, where substitution rates were allowed to vary freely amongst each branch, was significantly the best fit of all models, indicating that additional variation in substitution rates exists that cannot be explained by the differences in OGT alone.

### 3.4 Increased purifying selection in thermophilic species

Similar to seen in nucleotide substitution rates, dN/dS local rate models fit the data significantly better than the global rate model, regardless of whether a two-way split or three-way split in OGT classification was used (Table S2). Specifically, the dN/dS ratio was higher in branches delineating non-thermophilic species (0.064) than those delineating thermophilic species (0.059; Table 3). The three-way split model gave a statistically better fit than the global model or a local model that only distinguished thermophilic and non-thermophilic species. Thermotolerant taxa showed similar dN/dS values as thermophilic taxa (Table 3). The free-ratio model, where dN/dS rates are allowed to vary freely amongst each branch, was significantly the best fit of all tested models.

**Table 3:**
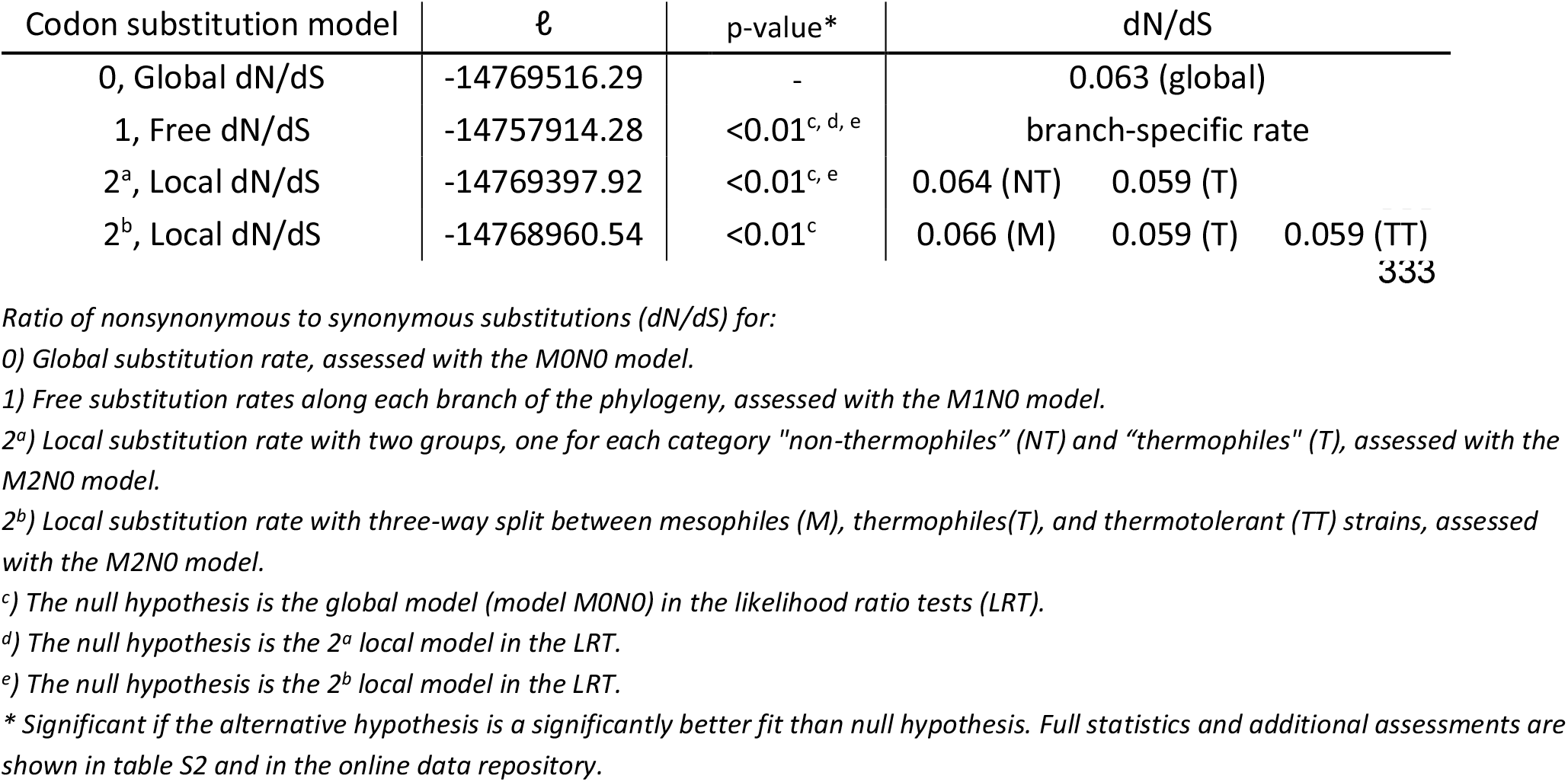
Summary statistics and parameter estimates of analyses of dN/dS among branches in the *Chaetomiaceae*.

| Codon substitution model | $\ell$ | p-value* | dN/dS | | |
| --- | --- | --- | --- | --- | --- |
| 0, Global dN/dS | -14769516.29 | - | 0.063 (global) |  |  |
| 1, Free dN/dS | -14757914.28 | <0.01<sup>c, d, e</sup> | branch-specific rate |  |  |
| 2<sup>a</sup>, Local dN/dS | -14769397.92 | <0.01<sup>c, e</sup> | 0.064 (NT) | 0.059 (T) |  |
| 2<sup>b</sup>, Local dN/dS | -14768960.54 | <0.01<sup>c</sup> | 0.066 (M) | 0.059 (T) | 0.059 (TT) |

Ratio of nonsynonymous to synonymous substitutions (dN/dS) for:
0) Global substitution rate, assessed with the M0N0 model.
1) Free substitution rates along each branch of the phylogeny, assessed with the M1N0 model.
2<sup>a</sup>) Local substitution rate with two groups, one for each category "non-thermophiles" (NT) and "thermophiles" (T), assessed with the M2N0 model.
2<sup>b</sup>) Local substitution rate with three-way split between mesophiles (M), thermophiles(T), and thermotolerant (TT) strains, assessed with the M2N0 model.
<sup>c</sup>) The null hypothesis is the global model (model M0N0) in the likelihood ratio tests (LRT).
<sup>d</sup>) The null hypothesis is the 2<sup>a</sup> local model in the LRT.
<sup>e</sup>) The null hypothesis is the 2<sup>b</sup> local model in the LRT.
\* Significant if the alternative hypothesis is a significantly better fit than null hypothesis. Full statistics and additional assessments are shown in table S2 and in the online data repository.

Both nonsynonymous and synonymous substitution rates were elevated in thermotolerant and thermophilic fungi compared to mesophiles. The mean nonsynonymous (dN) and synonymous (dS) substitution rates were 0.017 and 0.256 in mesophilic taxa, 0.019 and 0.323 in thermotolerant taxa, and 0.031 and 0.518 in thermophilic taxa (Table 4). Despite similar dN/dS ratios between thermotolerant and thermophilic species, thermophilic fungi exhibited substantially higher substitution rates. Thermophiles showed approximately 1.6-fold higher dN and dS values than thermotolerant species, and a stronger relative increase in dS (2.02-fold) than in dN (1.82-fold) compared to mesophiles (Table 4).

**Table 4:**
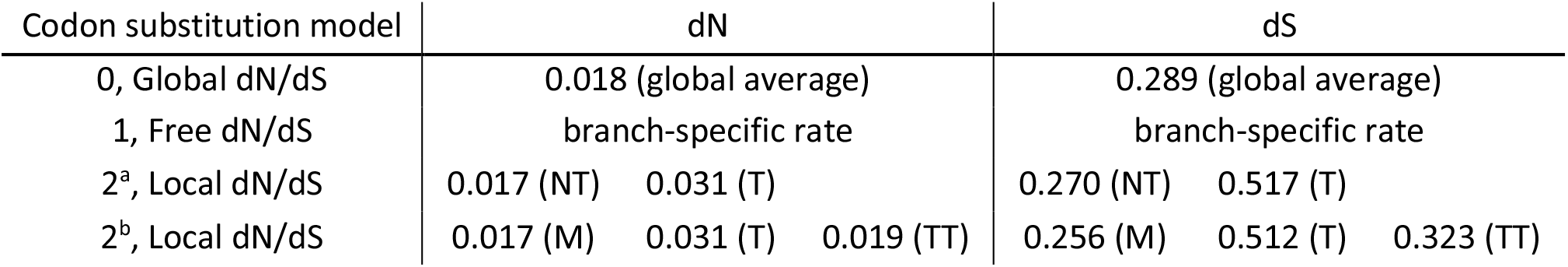
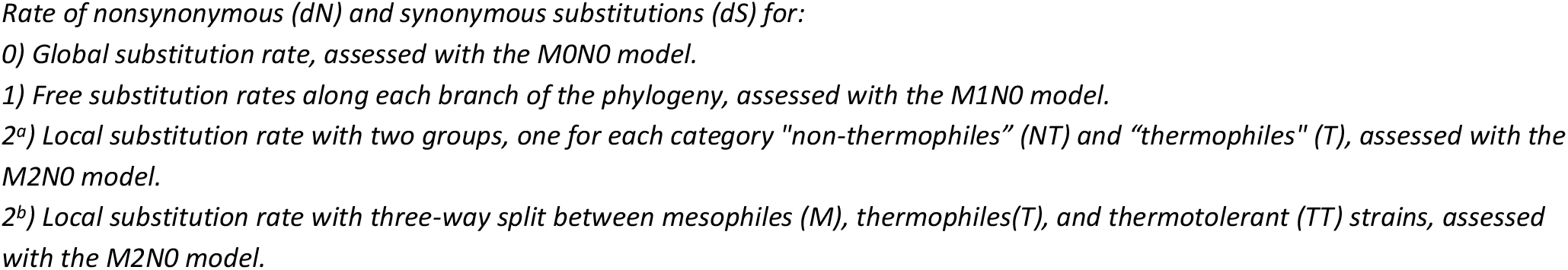
rates of nonsynonymous (dN) and synonymous (dS) substitutions amongst branches in the *Chaetomiaceae*.

| Codon substitution model | dN |  |  | dS |  |  |
| --- | --- | --- | --- | --- | --- | --- |
| 0, Global dN/dS | 0.018 (global average) |  |  | 0.289 (global average) |  |  |
| 1, Free dN/dS | branch-specific rate |  |  | branch-specific rate |  |  |
| 2<sup>a</sup>, Local dN/dS | 0.017 (NT) | 0.031 (T) |  | 0.270 (NT) | 0.517 (T) |  |
| 2<sup>b</sup>, Local dN/dS | 0.017 (M) | 0.031 (T) | 0.019 (TT) | 0.256 (M) | 0.512 (T) | 0.323 (TT) |

Rate of nonsynonymous (*dN*) and synonymous substitutions (*dS*) for: 0) Global substitution rate, assessed with the MONO model. 1) Free substitution rates along each branch of the phylogeny, assessed with the M1N0 model. 2<sup>a</sup>) Local substitution rate with two groups, one for each category "non-thermophiles" (NT) and "thermophiles" (T), assessed with the M2N0 model. 2<sup>b</sup>) Local substitution rate with three-way split between mesophiles (M), thermophiles(T), and thermotolerant (TT) strains, assessed with the M2N0 model.

Overall, these results show that thermophilic and thermotolerant species have increased rates of both nonsynonymous and synonymous substitutions on conserved orthologs, as well as stronger strength of purifying selection compared to mesophiles.

### 3.5 Optimal growth temperature is positively correlated with growth rate

To test the potential effect of growth rate on substitution rates, we subset the data to only include strains of the 19 species for which we measured growth rates. We assumed linear growth throughout the six days, the growth distance for the species tested at three days was doubled to estimate the growth distance after six days. We found a high correlation between growth rate and OGT (Pearson, rho = 0.62, p < 0.01, Figure 2). Mesophilic species grew slowest on average, with a growth of 46 ± 25 mm, thermotolerant fungi had average linear growth rates of 70 ± 25 mm, and the two thermophilic fungi grew fastest with average linear growth rates of 157 ± 4 mm. To account for the potential effect of non-linear mycelial growth, we performed an additional analysis that used a maximum growth of 85 mm within six days, which showed similar results (Table S1, Table S3).

**Figure 2:**
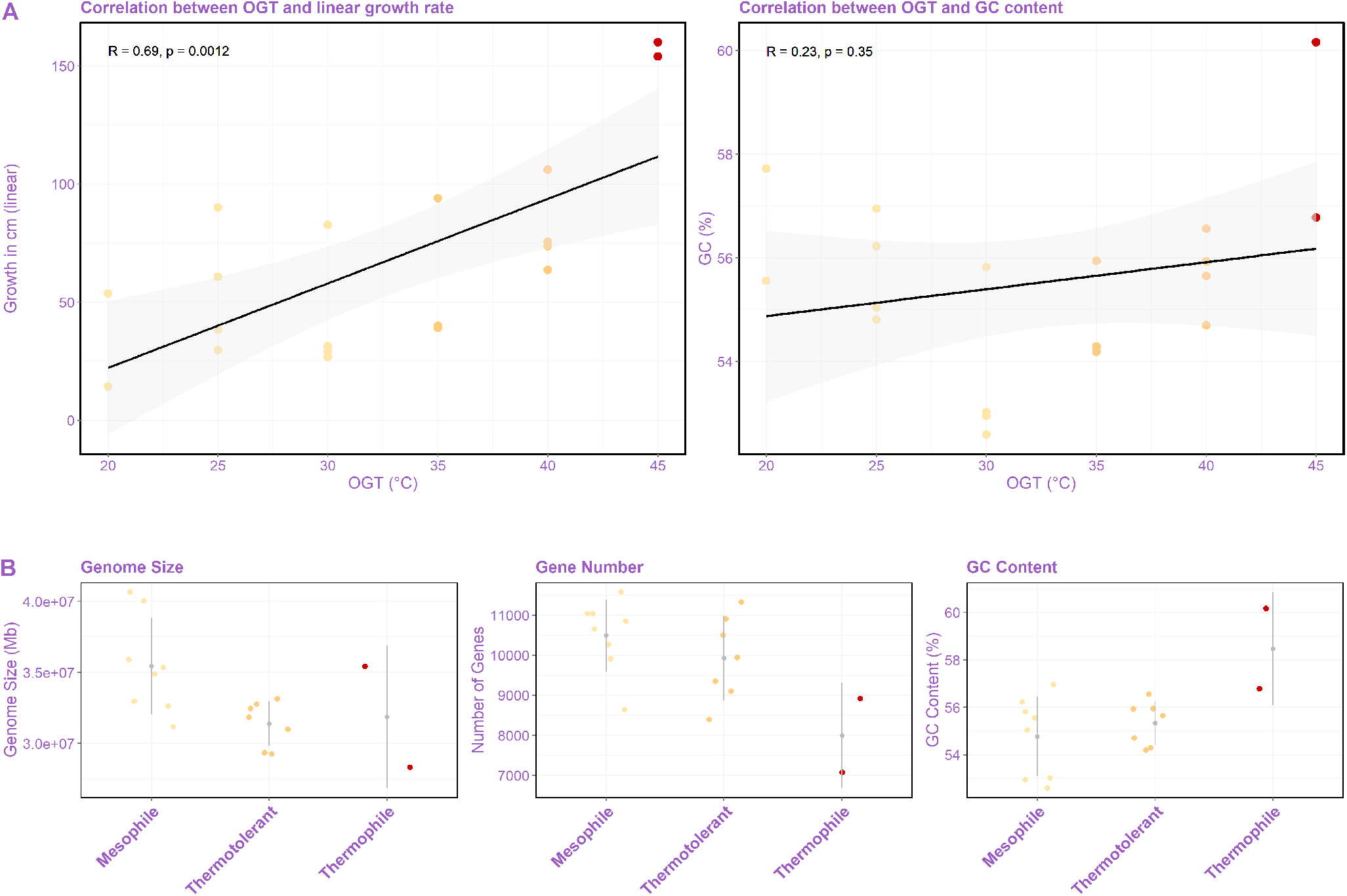
Correlation between optimal growth temperature and other variables. A) Pearson correlation coefficient (solid line) and 95% confidence intervals (grey-shaded area) are shown. Correlation between OGT and linear growth rate (left). Correlation between OGT and genome-wide GC content (right). B) Genomic differences between mesophilic, thermotolerant and thermophilic taxa. With from left to right genome size distribution, gene number distribution and genome wide GC content distribution.

### 3.6 Genomic features of thermophilic fungi

To explore the genomic features associated with a thermophilic lifestyle, we analyzed the effect of OGT and growth rate on several genomic variables, including effective number of codons (ENC), genome size, gene number, whole-genome GC content, GC content of conserved coding sequences, and GC content at third codon positions of conserved genes. OGTs were assessed in 5°C intervals, and Pearson correlation tests were used to assess relationships between OGT and each genomic trait (Table S1, Table S3).

In our dataset, codon bias levels (assessed with the effective number of codons) largely overlapped amongst genomes belonging to *Chaetomiaceae*. Pearson correlation coefficient showed no correlation between effective number of codons and OGT (R = 0.0, p-value = 0.97), and the effective number of codons was highly similar amongst mesophilic (ENC = 47.6 ± 3.7), thermotolerant (ENC = 48.6 ± 3.5) and thermophilic (average ENC = 48.8 ± 5.3) taxa.

Genome size showed an overall negative correlation with OGT (R = –0.57, p = 0.01). Genome size was highest in mesophilic taxa (37.3 ± 5.2 Mb), but thermotolerant (31.4 ± 1.6 Mb) and thermophilic taxa (31.9 ± 5.0 Mb) had similar genome sizes. Mesophilic taxa had higher genome sizes than all taxa with OGT ≥ 35°C (Figure 2). Gene numbers also decreased with increasing OGT (R=-0.62, p < 0.01), but the number of genes decreased more consistently along increasing OGT than genome size. Mesophilic taxa had the highest gene counts (11221 ± 1736), followed by thermotolerant (9930 ±1055) and thermophilic taxa (7797 ± 1319; Figure 2).

GC content, GC content of conserved coding sequences, and GC3 content exhibited a non-linear relationship with OGT: average GC content declined as OGT increased to ∼30°C, then rose again at higher temperatures (Figure 2; Table S1). On average, genome-wide GC content was comparable between mesophilic (55.1 ± 1.8 %) and thermotolerant taxa (55.3 ± 0.9%), but higher in thermophilic taxa (58.5 ± 2.4%). GC content of conserved coding sequences was largely similar between mesophilic (58.6 ± 1.9%), thermotolerant (59.1 ± 1.4%), and thermophilic taxa (61.2 ± 2.0%). A larger difference was observed in GC3 content, which ranged from an average of 74.9 ± 5.0% in mesophiles, and 76.1 ± 3.8% in thermotolerant taxa, to 82.0 ± 5.1% in thermophiles.

## 4. Discussion

Descriptions of OGT in *Chaetomiaceae* fungi have previously remained inconsistent as research has shown different OGT for the same fungal strains. Our dataset contained significant overlap with OGT data earlier obtained by Steindorff et al. (2024), and by van den Brink et al. (2015). Both of these studies used different laboratory methods and slightly different definitions of fungal thermophilia. We here unified the laboratory methods for the entire dataset, which resolved most of the differences initially observed between previous studies.

Nevertheless, despite similar laboratory methods, a few inconsistencies remained. Specifically, between van den Brink et al. (2015) and our study, two inconsistencies were found. *Achaetomium strumarium* and *Mycothermus thermophilus* were both classified as thermotolerant species in our study, while being classified as thermophilic in van den Brink et al. (2015). To this end, we note that *M. thermophilus* was capable of growing well at 45°C, despite showing an optimal growth at 40°C in our study. *A. strumarium* showed fast growth, and overgrew the entire plate within a six-day timespan. In redoing the analysis with a three-day incubation period, *A. strumarium* recovered an optimal growth at 35°C, and growth at 45°C was largely diminished. Furthermore, studies by Steindorff et al. (2024) recovered a number of different OGT than either our laboratory assessment or earlier accounts by van den Brink et al. (2015). Most notably, two species (*Chaetomidium leptoderma* and *Staphylotrichum longicolleum*) were identified as mesophilic in this study but described as thermophilic by Steindorff et al. (2024). Neither of these species were analyzed by van den Brink et al. (2015).

The exact reason for the discrepancies in OGT between the three studies cannot be assessed with the current methods, although we have two current hypotheses for the remaining discrepancies. Firstly, growth was assessed at the end of a three- or six-day incubation period in our study, while van den Brink et al., measured colony diameter before the colony at OGT had reached the edge of the Petri dish. Steindorff et al. (2015) followed the methodology earlier described by Morgenstern et al., (2012), which grew cultures from one to 32 days until differential growth amongst temperatures was clearly visible. Although our study standardizes laboratory methods by utilizing a set incubation time of six or three days, depending on growth rate, the caveat of this method is that it does not account for non-linear mycelial growth. The mycelial growth is normally characterized by a lag-phase after inoculation, followed by an exponential phase. After the exponential phase, a linear growth phase starts (see e.g. (Montini et al., 2006)). Not only the mycelial growth rate, but also the lag time can be temperature dependent (Negrão et al., 2021), which may have had an effect on the OGT assessed in this study. Beyond differences in laboratory methods, such large differences in OGT for both *C. leptoderma* and *S. longicolleum* could potentially also be explained by acclimatation to lower OGT in these laboratory strains, as the investigated strains of both these species have undergone a long laboratory history. Indeed, the strain of *C. leptoderma* (CBS 538.74) was isolated from English soil in 1974, while *S. longicolleum* (CBS 103.79) was isolated from pine vole dung in North Carolina in 1979. At the Westerdijk institute, the conditions for growth for *C. leptoderma* and *S. longicolleum* were described as 18°C and 24°C on cornmeal agar, respectively. In our assays, both species were indeed found to grow well at 20°C. As a result, we performed inoculations to other temperatures from stock plates kept at 20°C. Although not assessed in *Chaetomiaceae* fungi, other filamentous species such as *Metarhizium anisopliae* have been shown to overcome temperature barriers quickly (de Crecy et al., 2009). It would thus be possible that both species have undergone an adaptational shift to a lower OGT during our laboratory assessments. We believe that temperature adaptation experiments in this group of fungi are an important next step to further unravel adaptation to high temperature environments and could help disentangle the reasons behind the difference in assessed OGT between studies.

### 4.1 Rates of nucleotide substitution

Research suggests that high temperatures put different evolutionary pressures on different groups of organisms (Swami, 2009), and the mutagenic effect of above-optimal temperatures has been demonstrated in eukaryotes and prokaryotes alike (Chu et al., 2018; Matsuba et al., 2013). Plants and mammals from warmer environments tend to show higher substitution rates than those from more temperate regions (Gillman et al., 2009; Wright et al., 2006). For example, the substitution rate in the mitochondrial cytochrome b gene is on average 1.47 times higher in mammals living in warm climates than in cooler climates, while no difference is found in selection strength (Gillman et al., 2009). In thermophilic prokaryotes, on the other hand, research has shown a negative relationship between OGT and rates of evolution (Drake, 2009; Jacobs & Grogan, 1997; Mackwan et al., 2008). Thermophilic prokaryotes show a stronger selection against amino acid replacements (Friedman et al., 2004; Sabath et al., 2013), and a lower rate of genomic mutations than mesophiles (Drake, 2009). It has therefore been suggested that thermophilic prokaryotes, in contrast to mammals, have adapted to high temperatures by evolution of low substitution rates through e.g. optimization of DNA proofreading and repair mechanisms (Kontopoulos et al., 2020).

We here show that thermophilic fungi exhibit elevated nucleotide and amino acid substitution rates (in line with results from mammals and plants), along with stronger purifying selection on amino acid changes (in line with findings from thermophilic prokaryotes). To this end, we note that thermophilic bacteria and fungi have distinct differences, and considerable overlap, in their ecological niches. On the one hand, prokaryotic thermophiles are able to withstand wider ranges of temperatures: prokaryotic organisms are broadly classified as psychrophiles (OGT < 15°C), mesophiles (15 to 45°C), thermophiles (> 45°C), and hyperthermophiles (> 80°C; (Farrell et al., 2025). In comparison, thermophilic fungi grow at relatively moderate temperatures. The maximum theoretical temperature limit for eukaryotes lies around 62°C, above which thermostable and functional cellular organellar membranes cannot be formed (Patel & Rawat, 2021). In congruence with prokaryotes, and in contrast to mammals, however, thermophilic fungal species have adapted to function optimally in high-temperature environments (≥ 45°C). The increased strength of purifying selection in thermophilic fungi is thus consistent with the need to maintain protein stability in high-temperature environments (Kontopoulos et al., 2020).

This discrepancy between high nucleotide and amino acid substitution rates as well as high levels of purifying selection raises the question: why are nucleotide substitution rates higher in thermophilic fungi compared to mesophilic ones? One possibility is that thermophilic fungi experience elevated mutagenesis as a consequence of temperature-induced stress. If elevated temperature only increased mutation rates without altering selective constraints, synonymous and nonsynonymous substitution rates would be expected to increase proportionally, with little change in dN/dS. The lower dN/dS observed in thermophilic fungi instead suggests that elevated substitution rates are accompanied by stronger purifying selection against amino-acid-changing mutations. Thus, elevated mutagenesis cannot be ruled out, but it is unlikely to be the sole explanation for the observed pattern.

A more feasible explanation for this difference in nucleotide substitution rates is an increased substitution rate governed by a secondary factor, such as an elevated number of mitotic cell divisions per unit time (Nygren et al., 2011). Similar to results from prokaryotic thermophiles (Xu et al., 2021), growth rate and OGT were highly correlated in our dataset. A faster growth of thermophilic fungi than their mesophilic counterparts could lead to an elevated number of mitotic cell divisions per time unit in thermophilic fungi. As each replication carries the risk of incorporating the wrong nucleotides, this could consequently increase the substitution rate per unit of time in fast growing species compared to slow growing ones. Nevertheless, because little is known about the ecology of most *Chaetomiaceae* beyond their OGTs, we cannot rule out other explanations for the observed difference in substitution rates, such as difference in intrinsic mutation rates (Lynch, 2010), or generation time for the sexual life cycle (Nygren et al., 2011). We thus conclude that the higher rates of nucleotide substitution in thermophilic fungi may be caused directly by an increased mutagenesis of higher temperatures, indirectly by the increased growth rate of thermophilic fungi, or by other, currently unknown, factors.

### 4.3 Genomic properties and nucleotide composition

Beyond the rate of molecular evolution, multiple other genomic traits have been described to be influenced by high OGT. Thermophilic bacteria tend to have smaller genomes, which has been hypothesized to optimize energy utilization in stressful environments (Sabath et al., 2013). Indeed, we found here that both genome size and gene number decrease with increasing OGT in fungi, although the temperature at which effects were seen were trait dependent. Genome size was reduced in thermotolerant fungi compared to mesophiles, and did not decrease further in thermophilic fungi, whereas gene numbers show a more consistent decrease with increasing OGT. Together, these results suggest that genome streamlining in fungi might begin at OGT around 35°C.

An increase in GC content also correlates with a higher temperature optimum in bacteria (Hu et al., 2022; Musto et al., 2006; Wu et al., 2012; Yakovchuk, 2006), most likely because GC base pairs have increased thermal stability compared to AT base pairs (see e.g. (Yakovchuk, 2006)). Although thermophiles showed a higher genome-wide GC content than mesophilic or thermotolerant fungi, genome-wide GC content decreased on average up to an OGT of 30°C, indicating that GC-content increase could become increasingly important at higher growth temperatures, while being negligible at temperatures below 30°C. Indeed, similar correlations have been found in bacteria and archaea, where the correlation between tRNA GC content and OGT only becomes strongly positive at OGT above +-50°C (Hu et al., 2022). It is therefore likely that nucleotide content distributions would mainly play a role at higher temperatures than those assessed herein, or only become of importance at OGT beyond those of eukaryotic limits.

### 4.4 Conclusion

The *Chaetomiaceae* family contains a large number of thermophilic fungal species, which are interesting to research for the existence of thermostable enzymes, and opportunistic pathogens such as *Madurella mycetomatis* (Van Belkum et al., 2013; van den Brink et al., 2015). In this study, we show that OGT assessments of *Chaetomiaceae* fungi are not always straightforward. However, by standardizing the laboratory assessment, we confirmed existence of 2 thermophilic fungi within the family, in addition to the 9 already described by van den Brink et al., (2015). After refining the phylogeny of *Chaetomiaceae*, we found that thermophilic taxa show a lower dN/dS ratio, suggesting stronger purifying selection compared to mesophilic taxa. At the same time, thermophilic taxa showed higher nucleotide and amino acid substitution rates compared to mesophiles. This pattern suggests that thermophilic lineages accumulate more substitutions overall, but that amino-acid-changing substitutions are more strongly filtered, consistent with stronger purifying selection on conserved orthologs. The elevated substitution rates observed in thermophilic taxa may be partly explained by faster growth and a higher number of mitotic divisions per unit time, as OGT was positively correlated with growth rate. However, direct effects of temperature-induced mutagenesis, differences in life history, or other ecological and genomic factors cannot be ruled out.

Thermophilic fungal species showed lower genome sizes and gene numbers compared to mesophiles, and a higher genome-wide GC content. Together, our results strengthen knowledge on the correlation between fungal genomes and high OGT, by providing assessments of OGT and a phylogenetic framework, from which questions on the evolutionary consequences of fungal thermophilia were inferred.

## Data and materials

The computations/data handling were enabled by resources provided by the Swedish National Infrastructure for Computing (SNIC) at DARDELL, partially funded by the Swedish Research Council through grant agreement no. 2018-05973.

### Funding sources

We acknowledge funding from The Bergianus foundation/The Royal Swedish Academy of Sciences, and from the Swedish Research Council (grant 2024-04092) to Hanna Johannesson.

### Data availability

supplementary figures and - tables, together with gene alignments, result output files, and main scripts are available at the Zenodo repository: doi.org/10.5281/zenodo.21218879

